# Repurposing low–molecular-weight drugs against the main protease of severe acute respiratory syndrome coronavirus 2

**DOI:** 10.1101/2020.05.05.079848

**Authors:** Jia Gao, Liang Zhang, Xiaodan Liu, Fudong Li, Rongsheng Ma, Zhongliang Zhu, Jiahai Zhang, Jihui Wu, Yunyu Shi, Yueyin Pan, Yushu Ge, Ke Ruan

**Author notes:** J.G. and L.Z. contributed equally to this work.

## Abstract

The coronavirus disease (COVID-19) pandemic caused by infection with the severe acute respiratory syndrome coronavirus 2 (SARS-CoV-2) has affected the global healthcare system. Drug repurposing is a feasible method for emergency treatment. As low–molecular-weight drugs have high potential to completely match interactions with essential SARS-CoV-2 targets, we propose a strategy to identify such drugs using the fragment-based approach. Herein, using ligand- and protein-observed fragment screening approaches, we identified niacin and hit **1** binding to the catalytic pocket of the main protease of the SARS-CoV-2 (M^pro^), thereby modestly inhibiting the enzymatic activity of M^pro^. Chemical shift perturbations induced by niacin and hit **1** indicate a partial overlap of their binding sites, i.e., the catalytic pocket of M^pro^ may accommodate derivatives with large molecular sizes. Therefore, we searched for drugs containing niacin or hit **1** pharmacophores and identified carmofur, bendamustine, triclabendazole, and emedastine; these drugs are highly capable of inhibiting protease activity. Our study demonstrates that the fragment-based approach is a feasible strategy for identifying low–molecular-weight drugs against the SARS-CoV-2 and other potential targets lacking specific drugs.

---

The coronavirus disease (COVID-19) pandemic caused by the severe acute respiratory syndrome coronavirus 2 (SARS-CoV-2)^1-3^ has so far affected >4 million people worldwide, with a mortality rate of approximately 7%. Main protease (M^pro^ or 3CL^pro^) is one of the most extensively studied targets of coronaviruses^4^. M^pro^ plays an essential role in the cleavage of viral RNA-translated virus polypeptide^5^ and recognizes at least 11 cleavage sites in replicase polyprotein 1ab, e.g., LQ↓SAG (↓denotes the cleavage site). Covalent inhibitors against the SARS-CoV-2 M^pro^ have recently demonstrated potency toward inhibiting viral replication in cellular assays^6,7^; this further underpins the druggability of M^pro^. However, these compounds remain in the early stages of preclinical studies, and the development of new drugs usually takes years. The lack of drugs targeting the SARS-CoV-2currently poses a threat to numerous COVID-19 patients.

The COVID-19 pandemic has implored the repurposing of oral drugs^8^. As most recently approved drugs have been designed and optimized for specific targets, they are unlikely to completely match interactions with the SARS-CoV-2 targets. Compared with 13550 potential drugs in the DrugBank database at various stages from preclinical studies through approval, the estimated number of drug-like compounds (molecular weight of ∼500 Da) is reportedly approximately 10^60^. Therefore, the possibility of uncovering a highly potent and specific drug against the SARS-CoV-2 is quite slim. Conversely, low–molecular-weight drugs with intermediate potency and high safety can be an alternative treatment against the SARS-CoV-2. The toxicity of many low–molecular-weight drugs has been well understood owing to long clinical trials. Furthermore, their low structural complexity adds odds to fully match the interactions with anti-SARS-CoV-2 targets; e.g., the chemical space of compounds with <11 non-hydrogen atoms is approximately 10^9^. This is the cornerstoneof fragment-based lead discovery^9^, and many of the compounds in the fragment library were indeed extracted from pharmacophores of the approved drugs. We therefore hypothesize that it is highly possible to identify a low–molecular-weight drug containing pharmacophores using fragment-based screening (FBS).

We therefore assessed 38 compounds, which are pharmacophores (substructures) of repurposing drugs proposed from virtual screening, from our fragment library^10-15^ against the SARS-CoV-2 target. As these candidates have been predicted to bind at a high affinity, their pharmacophores should bind as well, albeit at a weaker affinity. These weak binders can be readily identified using a nuclear magnetic resonance (NMR) fragment-based approach, thereby eliminating the distracting false positives in virtual screening. Using our NMR fragment-based approach^16-18^, we characterized weak target–ligand interactions using ligand-observed spectra, e.g., saturation transfer difference (STD) and WaterLOGSY. Three hits for the SARS-CoV-2 M^pro^ were identified (Figure 1a): niacin, hit **1**, and hit **2** (Figure 1b). The potency of these three hits was then evaluated using enzymatic activity assay of the SARS-CoV-2 M^pro^ (Figure 1c). Niacin and hit **1** moderately inhibited the cleavage of fluorescent-labeled polypeptide (FITC-AVLQSGFR-Lys(Dnp)-Lys-NH2) by the SARS-CoV-2 M^pro^.

**Figure 1.**
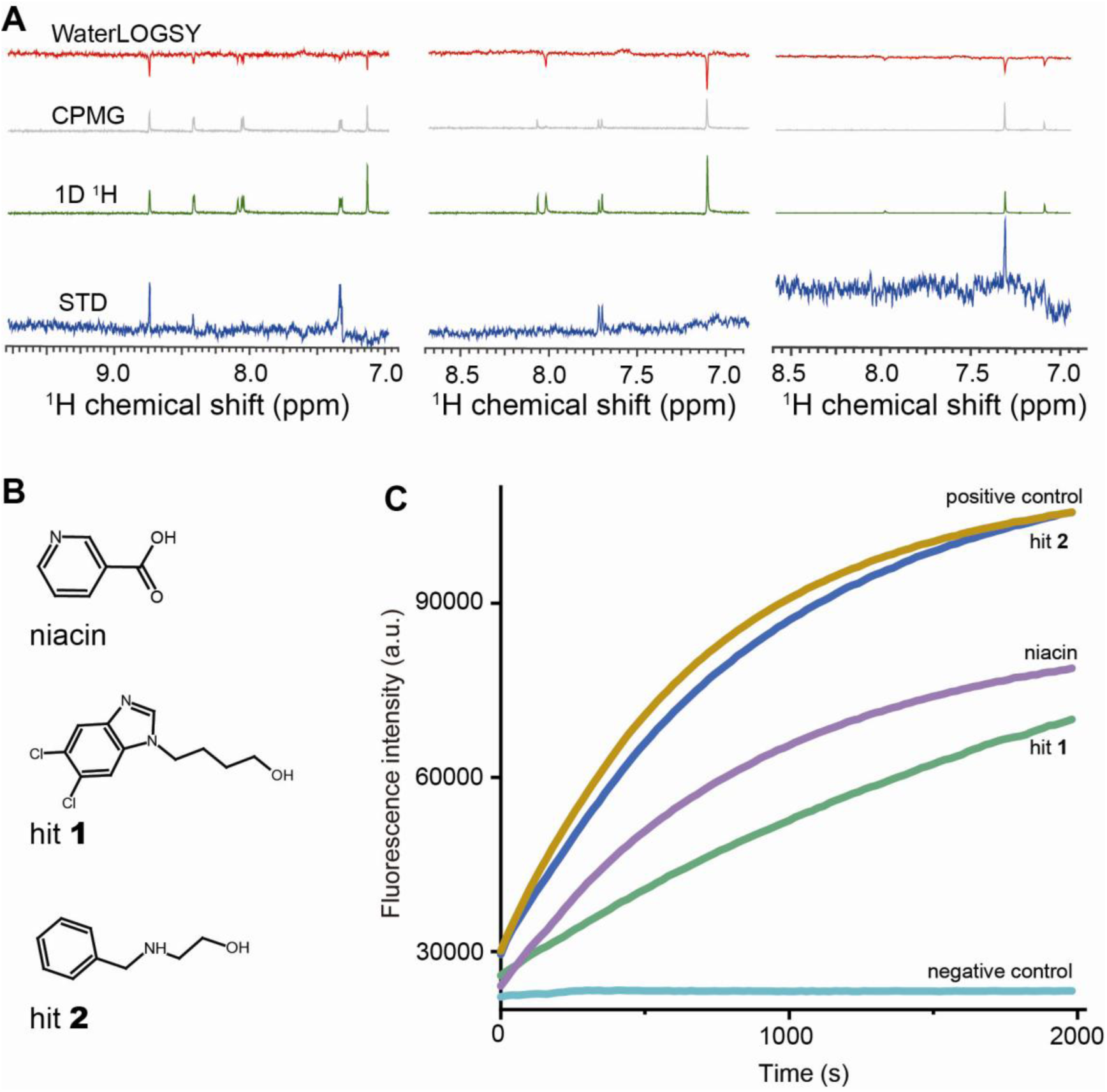
Fragment-based screening identified three hits of the SARS-CoV-2 main protease. **A**) NMR ligand-observed spectra of three hits in the presence of 10 μM full-length SARS-CoV-2 M^pro^. **B**) Chemical structures of the three hits. **C**) Inhibition of the enzymatic activity of the SARS-CoV-2 M^pro^ (5 μM) by the three hits (4 mM). The negative control was treated using fluorescence-labeled peptide (16μM) in the absence of M^pro^.

To further map the binding sites of niacin and hit **1**, we determined the chemical shift perturbations (CSPs) of ^15^N-labeled M^pro^ induced by the titration of these two compounds. Due to the severe signal overlap in the heteronuclear single-quantum correlation (HSQC) spectrum of the ^15^N labeled full-length SARS-CoV-2 M^pro^ (residue 4-306), the N-terminal domain(M^pro^-N) with the catalytic core included(residues 4–199) was used instead for all the protein-observed NMR studies conducted thereafter. The well-dispersed HSQC spectrum of the SARS-CoV-2 M^pro^-N enabled many ^1^H-^15^N amide chemical shift assignments transferred directly from SARS-CoV M^pro^-N^19^, as these two proteins share 96% sequence identity. Key residues proximal to the catalytic site, including H41, V42, D48, N51,G143, H163, and V186, were thus assigned (Figure S1). Both niacin and hit **1** perturb a common residue V42, suggesting that these two hits bind to the catalytic core of SARS-CoV-2 M^pro^. Interestingly, these two hits also recognize different sets of residues in the catalytic core; for example, niacin perturbs H41, G143, N51, and V186 (Figure 2a), whereas hit **1** induces CSPs of residues M165, E166, and L167 (Figure 2b). Mapping of these residues to the surface representation of the crystal structure of the SARS-CoV-2 M^pro^ (PDB code: 6LU7)^6^ suggested that these two hits adopted different orientations in the catalytic site, with a shared anchor point near V42 (Figure 2c). Considering the molecular size of niacin and the spatial distribution of the perturbed residues, niacin probably binds more than one site. Nevertheless, the CSP pattern suggests that the catalytic core of the SARS-CoV-2 M^pro^ accommodates compounds larger than niacin and hit **1**.

**Figure 2.**
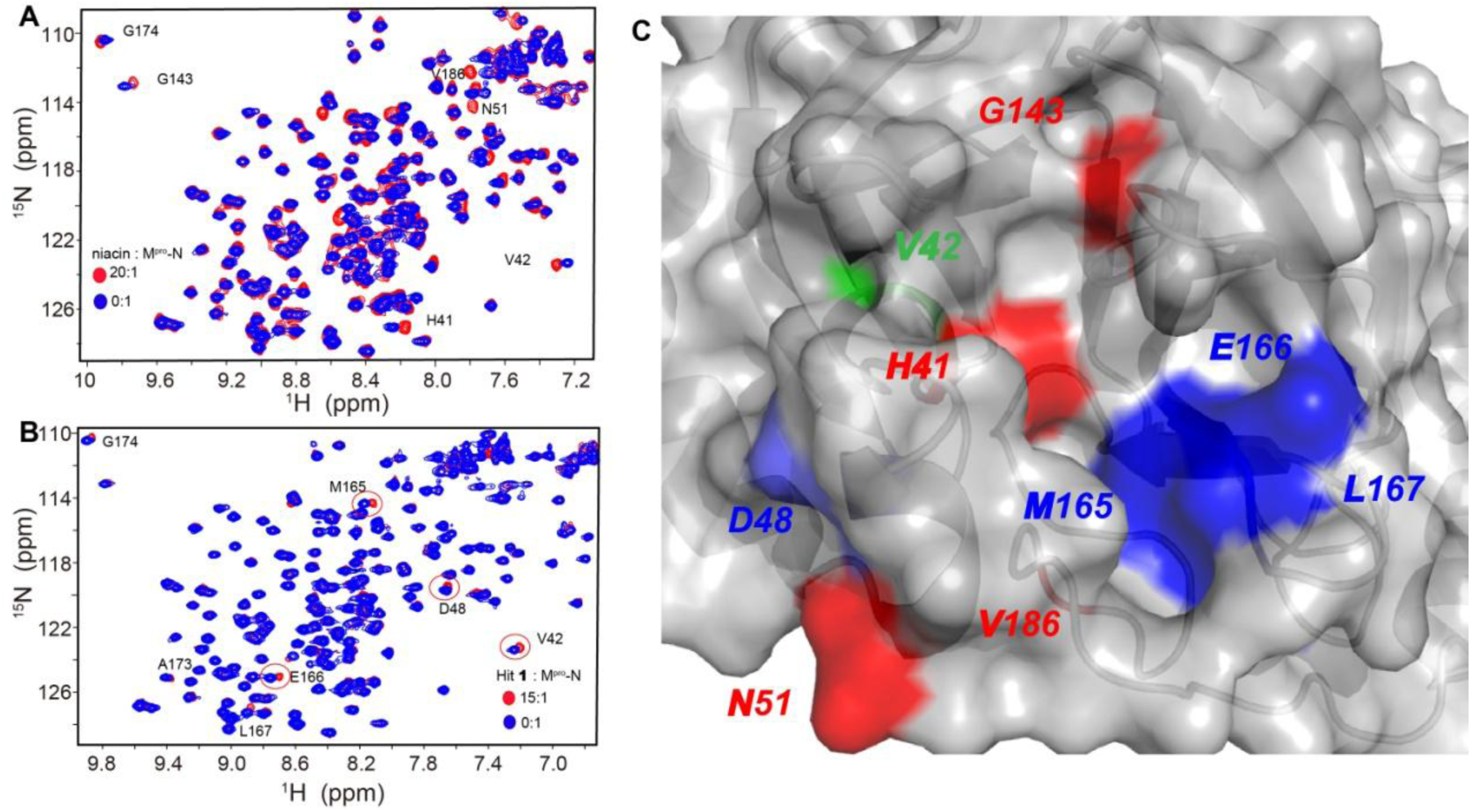
Binding topology of niacin and hit **1**determined from NMR chemical shift perturbations. **A**) Superimposition of 2D ^1^H-^15^N HSQC spectra of the SARS-CoV-2 M^pro^-N in the absence and presence of niacin. The ligand:protein molar ratios are shown. **B**) Chemical shift perturbations induced by hit **1. C**) Chemical shift perturbations induced by niacin (red), hit **1** (blue), or both (green) were mapped to the surface of the crystal structure of the SARS-CoV-2 M^pro^ (PDB code 6LU7).

We therefore searched for low–molecular-weight (<400 Da) drugs containing the pharmacophores of niacin and hit **1**. Molecular docking identified carmofur as a niacin derivative. Titration of carmofur with ^15^N-labeled SARS-CoV-2 M^pro^-N induced an extra setof cross peaks at a ligand:protein molar ratio of 2:1, and some original signals completely disappeared at a molar ratio of 4:1 (Figure S2a). This indicated a strong binding between carmofur and SARS-CoV-2 M^pro^, as confirmed by the IC_50_ of 2.8 ± 0.2 µM determined using enzymatic assay at an M^pro^ concentration of 0.5 µM (Figure S2b). However, the original NMR signals completely disappeared at a ligand:protein molar ratio that significantly deviated from a stoichiometry of 1:1. It has been recently demonstrated that carmofur is a covalent inhibitor of the main protease of theSARS-CoV-2, with an IC_50_value of 1.82 μM in the presence of 0.2 µM enzyme^6^; this was consistent with our measurement as a higher M^pro^ concentration was used in our case. Collectively, these data suggest that covalent linking to the SARS-CoV-2 M^pro^ is driven by excess carmofur. Nevertheless, using this fragment-based approach is a feasible strategy for repurposing low–molecular-weight drugs targeting SARS-CoV-2 M^pro^.

Further pharmacophore identification and molecular docking nominated several low–molecular-weight analogs of hit **1**, for example, triclabendazole, emedastine, and bendamustine (Figure S3a). The single-dose enzymatic assay showed that these three drugs had significantly higher potency than hit **1** (Figure S3b). We further determined the dose-dependent response of bendamustine and emedastine in the inhibition of the SARS-CoV-2 M^pro^ activity, with IC_50_ values of 26 ± 1 μM and 82 ± 7 μM (Figure 3a). The IC_50_ value of triclabendazole was roughly estimated to be 70 μM from the two-dose inhibition rates (31% and 72% inhibition at 50 μM and 100 μM triclabendazole, respectively), as limited by the low aqueous solubility of triclabendazole. Further, bendamustine and emedastine induced significantly larger CSPs in a dose-dependent manner than hit **1** (Figure 3b).

**Figure 3.**
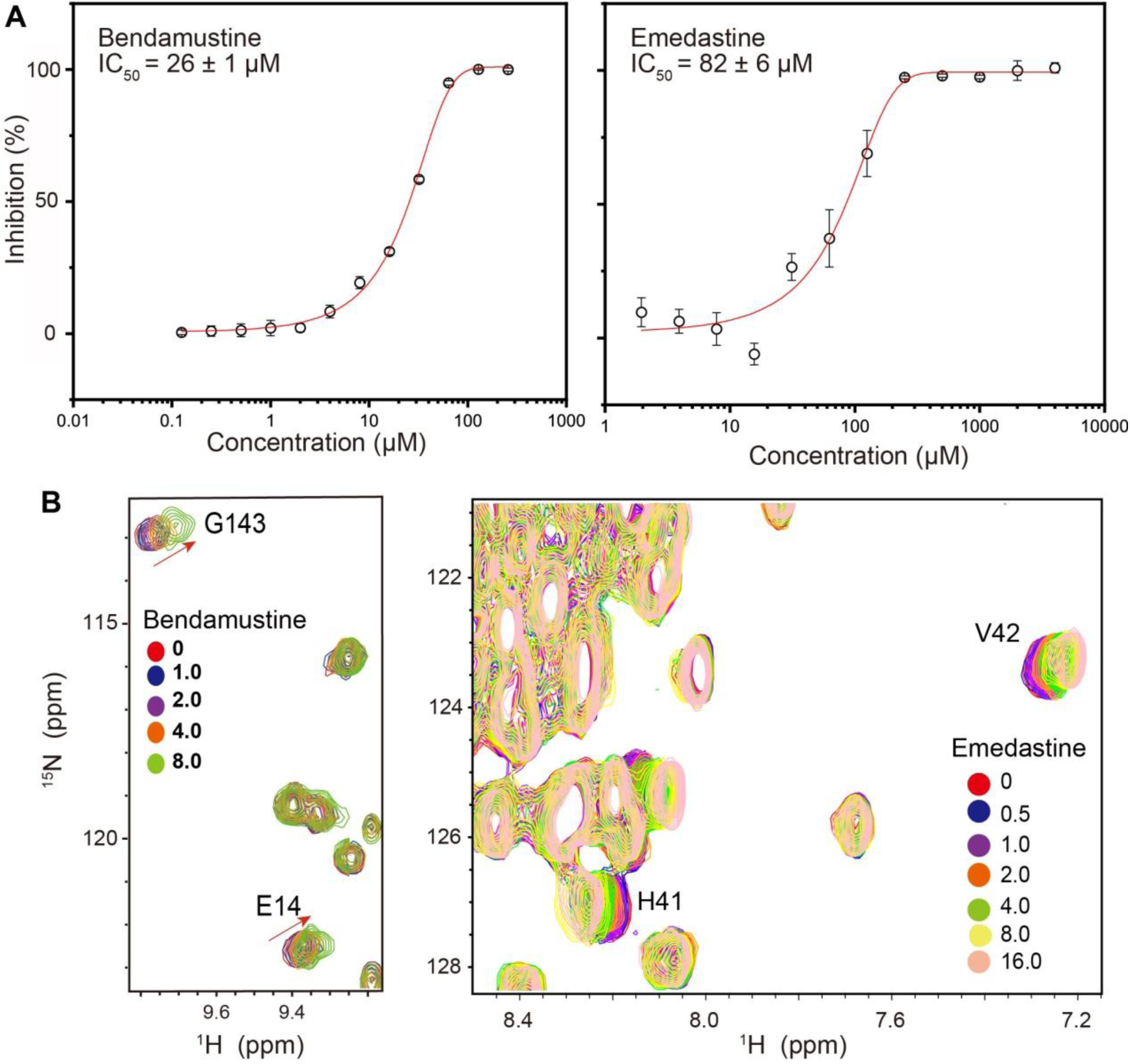
Potency and binding topology of low–molecular-weight drugs as derivatives of hit **1. A**) The dose-dependent inhibition of the enzymatic activity of the SARS-CoV-2 M^pro^ (0.5 μM) by bendamustine and emedastine in the presence of 16 μM fluorescent-labeled substrate. **B**) Chemical shift perturbations of ^15^N-labeled SARS-CoV-2 M^pro^-N induced by bendamustine and emedastine at the annotated ligand:protein molar ratio.

Taken together, our fragment-based strategy facilitates the identification of low–molecular-weight drugs against the SARS-CoV-2 M^pro^. First, this approachcan be readily applied to identify low–molecular-weight drugs against other SARS-CoV-2 targets (e.g., RNA-dependent RNA polymerase or the receptor-binding domain of the spike protein). Second, a combination of these low–molecular-weight drugs may be used to gain higher potency than that achieved via a single compound if their binding topologies show no evidence of steric repulsion. Finally, although carmofur and bendamustine show higher potency than the fragment screening hits, the toxicity of anti-cancer drugs remains a challenge in their clinical applications. Conversely, triclabendazole and emedastine could be valuable in inhibiting the SARS-CoV-2 replication at the early stage. It may also serve as a starting point for the next round of pharmacophore identification. In general, our study provides new insights toward the repurposing of low–molecular-weight drugs against the SARS-CoV-2 and other potential targets lacking specific drugs.

## ACKNOWLEDGMENT

Part of our nuclear magnetic resonance study was performed at the National Center for Protein Science Shanghai andthe High Magnetic Field Laboratory of the Chinese Academy of Sciences. We thank the Ministry of Science and Technology of China (2016YFA0500700 and 2014CB910604), the National Natural Science Foundation of China (21874123, 21703254 and 21807095), the Fundamental Research Funds for the Central Universities (WK2060190086), for the financial support, and supercomputing resources from the Bioinformatics Center of the University of Science and Technology of China, School of Life Sciences.

