## Supporting Information for "Repurposing low–molecular-weight drugs against the main protease of severe acute respiratory syndrome coronavirus 2" (from another version of this preprint: version 1, 2020-05-06)

### Materials and Methods

*Protein expression and purification.*DNA encoding human$M^{pro}$（residues 1-306）was amplified and incorporated into a modified pGEX-4T-1 (Novagen) plasmid, in which the thrombin protease site was substituted for a tobacco etch virus (TEV) cleavage site. The proteins were expressed in *Escherichia coli* BL21(DE3)-RIL cells (Novagen) cultured in LB mediumat 37℃ till $OD_{600}$ reaches about 0.8-1.2, then shifted to 16℃and induced with 0.4 mM isopropyl-β-D-thiogalactopyranoside (IPTG)for 24 hours. Bacterial pellets were resuspended in buffer A (20 mM Tris, 1 M NaCl, 2 mM DTT, 1 mM EDTA, pH 7.6) and lysed by sonication on ice. TheGST-tagged proteins were purified on glutathione-Sepharose beads (GE Healthcare)，then treated with TEV to cleave the N-terminal GST tag overnight at 16℃.The protein was then purified by a Superdex 200 column (GE Healthcare).Finally, the purified full-length $M^{pro}$ proteins（residues 1-306）were concentrated to ~10 mg/ml in buffer B (0 mM Tris, 150 mM NaCl, 2 mM DTT, 1 mM EDTA, pH 7.6) and stored at -80℃ for 1D NMR screening, crystallographic study orenzymatic assay.

The N-terminal Mpro proteins（residues 4-199）were mainly used for NMR research.DNA encoding N-terminal Mpro proteins (residues 4-199) was cloned into the pET22b vector(GE Healthcare,Shanghai,China).Thereconstruct plasmids were transformed to Escherichia coli BL21(DE3)-RIL. Cells were grown in minimal medium supplemented with 15NH4Cl for NMR samples. Generally, the proteins expression were induced at OD600 of 0.8-1.2using 0.4mM IPTG at 16℃ for 24 hours. All proteins were purified using Ni-chelating column(Qiagen, GE Healthcare) and further purified using gel filtration chromatopraphy(superdex-200 16/60).

*NMR fragment-based screening.* We applied a highly automated method of NMR fragment based screening to discovered 3 hits of $M^{pro}$(residues 1-306 )protein from 38 compounds in our fragment library. The NMR screening samples were parpered containing 0.4mM compound in DMSO-d6 solution,50% D2O and 10uM unlabeled $M^{pro}$ protein. The binding hits were identified based on the signals of STD and WaterLOGSY. The whole process of NMR fragment-based screening was performed at 298k on a 700 MHz spectrometer (Agilent Technologies Santa Clara, CA, USA) equipped with a 96-well autosampler and a cryoprobe.

*Chemicalshift perturbations.* The NMR HSQC titration was applied to detect the residues perturbed by chemicals and determine the affinity of N-terminal Mpro and compounds.The HSQC spectrums were collected for each titration points at Agilent 700MHz spectrometer equipped with a cryoprobe, with a total experimental time of 71 minutes at 308k. The compounds were stocked in DMSO at a concentration of 20 mM.The 0.1mM 15N-labeled MPRO (residues 4-199) proteins were titrated at the ligand/protein molar ratio ranging from 0.0 to 8.0,respectively.

*Enzymatic assay of SARS-CoV-2 $M^{pro}$.*To characterize the enzymatic activity of SARS-CoV-2 $M^{pro}$ and to measure the inhibition activities of different inhibitors, the fluorescence resonance energy transfer (FRET) based assay which reported before was carried out [ref: Yang H, Xie W, Xue X, Yang K, Ma J, et al. (2005) *PLoS Biol* 3(10): e324.]. In brief, the enzymatic activity was measured by continuous kinetic assays using the fluorescently labeled substrate, FITC-AVLQSGFR-Lys(Dnp)-Lys-NH2 (over 95% purity, GL Biochem Shanghai Ltd, Shanghai, China).The fluorescence intensity was monitored with a CLARIO star microplate reader (BMG Labtech, Germany) using exciting and emitting wavelengths at 485nm and 520 nm, respectively. The experiments were performed with abuffer consisting of 50 mM Tris-HCl, 1 mM EDTA, 5% DMSO and the pH is 7.3.The SARS-CoV-2 $M^{pro}$ protease (final concentration of 0.5 μM~5 μM)was mixed with the inhibitors and incubated at 30℃ for 10 minutes. After centrifuge at

13000 rpm and 4℃ for 10 minutes, the supernatant of the mixture was added into the substrate, whose final concentration was 10 μM~16 μM and start the measurement immediately until up to 50 minutes with 20 or 30 seconds interval. The data was analyzed by OriginPro 2020 pro (OriginLab Corporation, Northampton, Massachusetts, USA).

*Molecular docking.*All structures of chemical compounds were obtained from DrugBank database. 3D structures of all small molecules were minimized with Tripos force field after adding Gasteiger and Marsili charges. Crystal structure of Mpro (PDB ID : 6LU7) was obtained from PDB database and prepared with the following instructions: All hydrogen atoms were added with H-bonding orientation;  All ligands and water were removed; repaired related missing sidechains were repaired and atom bumps were removed with 100 cycles energy minimized. Small molecules were docked with Mpro using Autodock Vina software. Docking site was defined as the cubic grid box with side length of 22 angstroms centered on the catalytic active site. For virtual screening, the binding modes with the lowest binding energy were recommended for each molecule.

**Figure Legends**

**Figure S1.** The backbone $^{1}H$-$^{15}N$ chemical shift assignment of SARS-CoV-2 $M^{pro}$-N transferred from its homolog SARS $M^{pro}$-N.

**Figure S2.** Carmofur binds to the catalytic core and inhibits the enzymatic activity of SARS-CoV-2 $M^{pro}$. **A**) Chemical shift changes induced by carmofur at various doses. B) Dose-dependent inhibition profile of carmofur in the presence of 0.5 μM SARS-CoV-2 $M^{pro}$ and 16 μM fluorescence labeled substrate. Errors were estimated from three biological duplicates.

**Figure S3.** Enzymatic activity of SARS-CoV-2 $M^{pro}$ inhibited by repurposing drugs containing hit 1 pharmacophore. **A**) The chemical structures of the drugs as hit **1** derivatives with molecular docking energy annotated. **B**) The single-dose (60 μM) inhibition of the enzymatic activity of SARS-CoV-2 $M^{pro}$ with inhibition rate annotated.

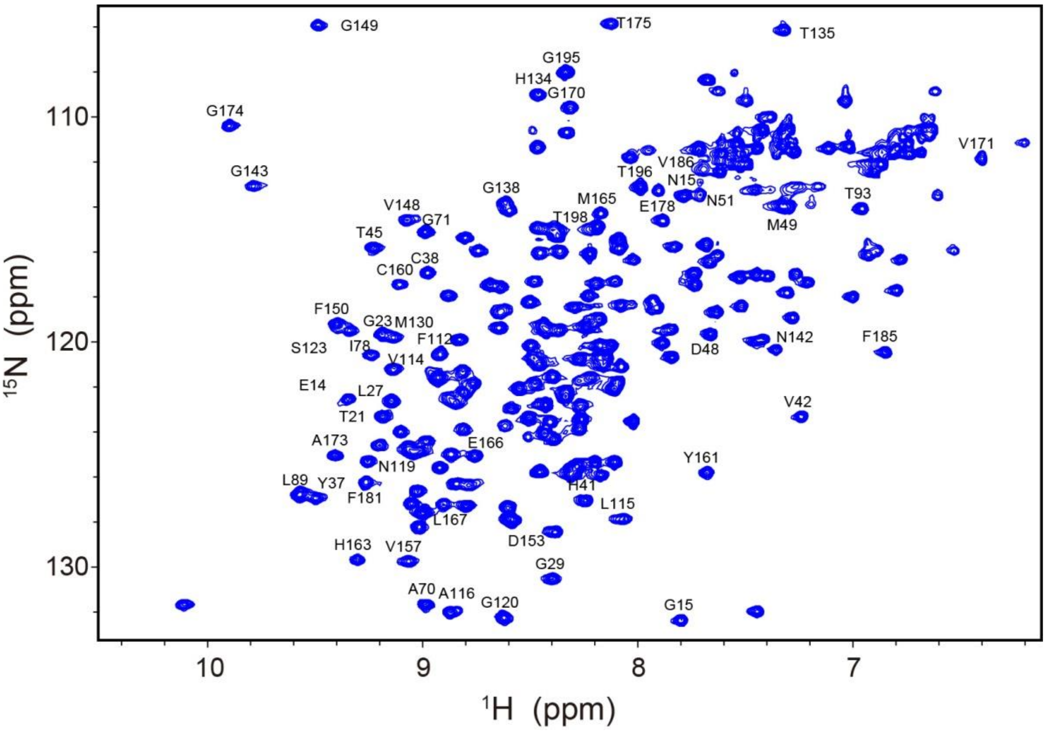

G149
T175
T135
G195
H134
G170
G174
V171
G143
T196
V186
N15
G138
M165
E178
N51
T93
V148
G71
T198
M49
T45
C38
C160
F150
G23
M130
F112
N142
F185
S123
I78
D48
V114
E14
L27
V42
T21
A173
E166
Y161
N119
L89
Y37
H41
F181
L115
L167
D153
H163
V157
G29
A70
A116
G120
G15
110
120
130
10
9
8
7
$^{15}$N (ppm)
$^{1}$H (ppm)


**Figure S1.**

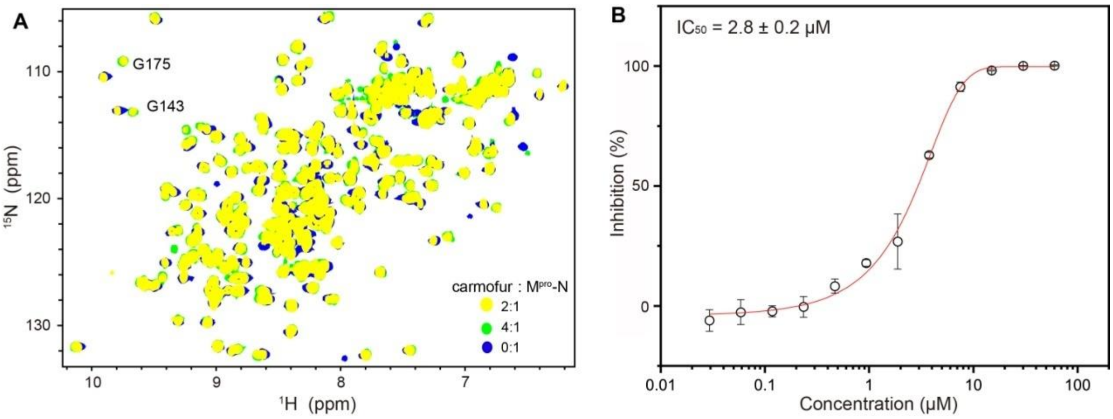
A
G175
G143
carmofur : Mpro-N
2:1
4:1
0:1
15N (ppm)
1H (ppm)
B
IC50 = 2.8 ± 0.2 μM
Inhibition (%)
Concentration (μM)

**Figure S2.**

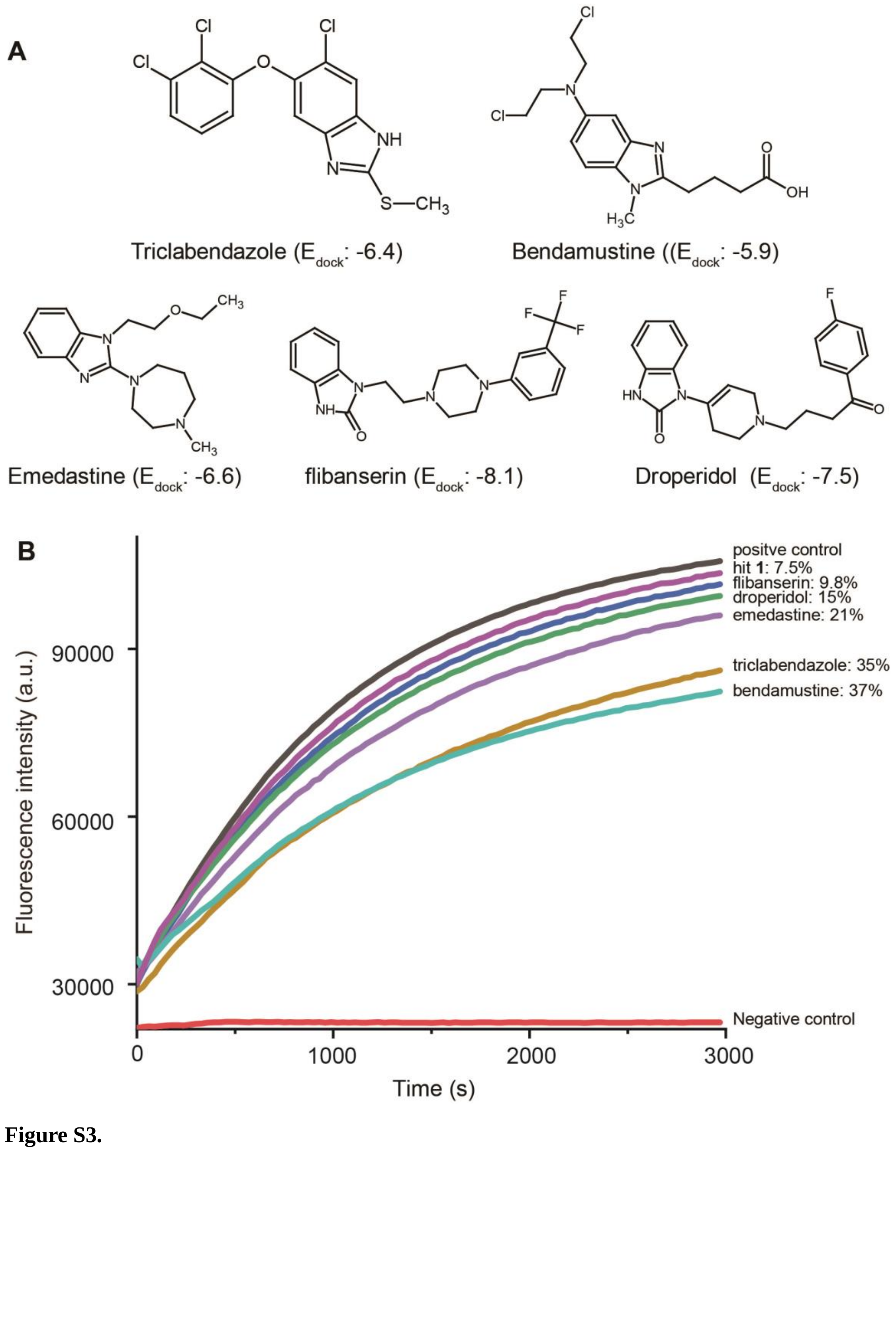
A
Triclabendazole ($E_{dock}$: -6.4)
Bendamustine (($E_{dock}$: -5.9)
Emedastine ($E_{dock}$: -6.6)
flibanserin ($E_{dock}$: -8.1)
Droperidol ($E_{dock}$: -7.5)
B
positve control
hit **1**: 7.5%
flibanserin: 9.8%
droperidol: 15%
emedastine: 21%
triclabendazole: 35%
bendamustine: 37%
Negative control
Fluorescence intensity (a.u.)
Time (s)

**Figure S3.**
